# FIDDL: depth-matched negative controls distinguish genuine interspecific introgression from competitive-mapping artifact

**DOI:** 10.64898/2026.08.06.743240

**Authors:** Kaitlyn Taylor, Kennadi A. Shumaker, Spencer J. Gray, Matthew L. Bochman

## Abstract

Interspecific introgression is routinely detected by competitively mapping reads to a concatenated multi-species reference and calling regions where a non-focal species recruits coverage. Using strains that cannot contain the ancestry being detected, we show this design generates substantial false-positive signal through two mechanisms with opposite phylogenetic-distance signatures. Standard nuclear assemblies omit the mitochondrion and 2-micron plasmid, leaving high-copy cytoplasmic reads without a legitimate target; completing the reference preferentially removes signal from the most divergent donor. Genuine cross-species sequence conservation inflates the most closely related donor. Masking chromosome ends removes its subtelomeric part but plateaus at a non-zero floor, and the interior residual traces to conserved single-copy genes where a short read carries under one base of discriminating information. The floor grows with sequencing depth (1.19% of callable positions at 50×, 2.02% at 147×, 3.85% at 393× in a pure strain), is not mitigated by long reads, and appears at sub-diploid dosage – three properties widely read as evidence of authenticity. Because the discriminating information is below single-read resolution, no read-level filter separates artifact from introgression; we show three that fail. What works is locus-level: a consensus-phylogenetic test (29/29 specificity on confirmed artifact) and an allele-fraction donor-match test, complementary and validated in both directions on independent published introgression. We package the comparative controls as FIDDL (*False Introgression Detection via Depth-matched controls and Loci-recurrence*), an open-source tool, withdraw two of our own analysis-ready calls, and show re-analysis of published wild isolates reduces low-confidence introgression by ∼53% while leaving high-confidence signal intact.

## INTRODUCTION

Interspecific gene flow is central to the evolutionary genomics of *Saccharomyces*. Hybridization produced the lager yeasts and a growing catalogue of natural hybrids, and introgressed tracts segregate widely in domesticated and wild *S. cerevisiae* (Dunn and Sherlock 2008; Libkind, et al. 2011; Morales and Dujon 2012; Peter, et al. 2018). The scale of these inferences has grown accordingly: introgression is now quantified across panels of hundreds to thousands of genomes, and the resulting counts feed downstream claims about adaptation, domestication history, and the permeability of species boundaries (Gallone, et al. 2016; Yue, et al. 2017; Peter, et al. 2018).

Most large-scale studies rely on a common methodological pattern. Reads are mapped competitively against a reference formed by concatenating the nuclear genomes of several species. Each read is then assigned to whichever species it aligns to best, and regions where a non-focal species accumulates coverage above a threshold are called introgressed (Dunn and Sherlock 2008; Yue, et al. 2017; Peter, et al. 2018). The approach is attractive because it is cheap, requires no phasing or assembly, and scales to large panels. It is also, as we show here, systematically biased in a direction that inflates precisely the quantity it is used to measure.

The vulnerability is easy to state. Competitive mapping forces every read to a decision, but the reference is not a complete representation of the genomes being compared. Standard nuclear assemblies, including the *Saccharomyces cerevisiae* S288C reference, omit the mitochondrial genome and the 2-micron plasmid (Goffeau, et al. 1996; Engel, et al. 2014), high-copy elements that can constitute a large fraction of a whole-genome library (Broach 1982; Foury, et al. 1998). Reads from those elements have no legitimate target in a nuclear-only reference and must map somewhere, be discarded, or map poorly (Li and Durbin 2009). Separately, and independently of reference completeness, subtelomeric regions of *Saccharomyces* chromosomes are enriched for gene families and repeat classes genuinely conserved across species (Louis 1995; Yue, et al. 2017). Reads from these regions align nearly as well to a congener as to the focal species, and competitive assignment resolves the near-tie arbitrarily.

Neither observation is new in principle. That organellar sequence should be included in a reference, and that subtelomeres are difficult, are both familiar cautions (Engel, et al. 2014). What has not been systematically evaluated is the magnitude of the resulting error, its dependence on sequencing parameters, whether it is mitigated by the long-read technologies now displacing short reads, whether the heuristics commonly used to validate introgression calls can distinguish it, and – most practically – what analysis a working researcher should actually run to determine whether their own results are affected.

Here, we measure the false-positive floor directly, decompose it into two mechanisms with distinguishable phylogenetic signatures, characterize its dependence on depth and library composition, and show that two widely used validation heuristics fail against it. We then package the controls that do work as FIDDL (*False Introgression Detection via Depth-matched controls and Loci-recurrence*), an open-source command-line tool. We validate FIDDL by applying it to two of our own introgression calls – both analysis-ready, both supported by conventional evidence – and withdrawing them, and by quantifying how much apparent introgression in a recent published panel is attributable to the same mechanisms.

## RESULTS

### Depth convention

Two depth conventions appear below and are not interchangeable. Short-read depths are bp-weighted means over focal-species nuclear windows, matching the coverage-ladder computation. Long-read depths are medians over focal-species nuclear windows, matching the convention of the original long-read analysis. For the short-read control, these differ by 13-15% (*e.g.*, 147.13× mean *vs.* 127.18× median for the same condition), so values should not be compared across platforms without conversion.

### A false-positive floor that grows with sequencing depth

*S. cerevisiae* S288C carries no *Saccharomyces eubayanus* or *Saccharomyces paradoxus* ancestry (Engel, et al. 2014; Peter, et al. 2018). Therefore, any interspecific signal recovered from S288C reads is false by construction. We mapped a whole-genome Illumina library from S288C (SRA run SRR2070491; 42.9 M read pairs, 6.40 Gb) competitively against a reference comprising the S288C nuclear chromosomes together with the *S. eubayanus* (FM1318) and *S. paradoxus* (CBS432) nuclear chromosomes, and no organellar sequence from any species – hereafter referred to as REF-naive, replicating the standard published design.

The floor is large and rises continuously with sequencing depth (Fig. 1A). Down-sampling the control across a coverage ladder gave apparent non-cerevisiae ancestry of 0.026% at 10×, 0.109% at 25×, 1.19% at 50×, 1.69% at 100×, and 2.14% at 200×, without saturating; the library’s full depth (393×) gave 3.85%. No *S. eubayanus* windows were called at all below 50×. For comparison, published introgression fractions for individual wild S. cerevisiae isolates frequently fall between a few tenths of a percent and a few percent (Barbosa, et al. 2016; Peter, et al. 2018). In other words, at depths now routine, the false-positive floor of the naive design is of the same order as the biological signal it is used to detect.

**Figure 1.**
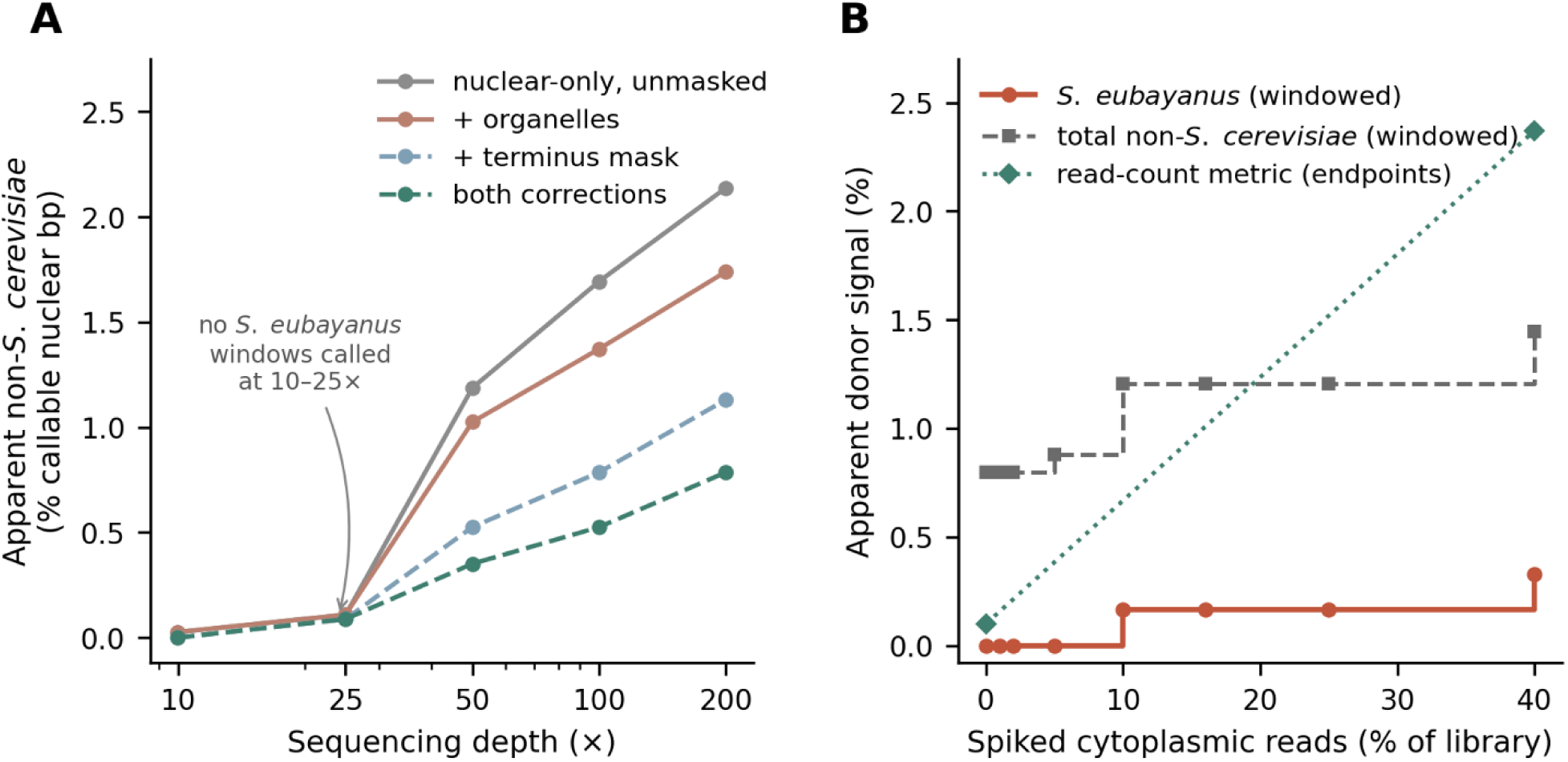
The artifact is a function of sequencing depth and library composition. **(A)** Apparent non-*cerevisiae* fraction *vs*. focal-species sequencing depth for the S288C negative control (10×, 25×, 50×, 100×, 200×), under both references and both masking conditions; no *S. eubayanus* windows are called at 10× or 25×. **(B)** Apparent donor signal *vs*. spiked cytoplasmic read fraction (0-40%) on a read-count metric. The windowed metric on the same data is shown as a step function for comparison.

Because the calling threshold is an absolute depth (≥5×), a mechanical explanation is available: as depth rises, artifactual windows cross a fixed bar, while the focal-species denominator is already saturated. We tested this by reclassifying at a threshold scaled to each library’s own median depth. The depth trend persisted essentially unchanged (−12.8% under a fixed threshold *vs*. −14.6% under a scaled threshold across the same depth interval), indicating that the depth dependence is a property of the mapping behavior rather than an artifact of thresholding.

The signal is not diffuse noise but resolves into discrete blocks. At 147×, thresholding retained 61, 41, 24, 21, and 9 *S. paradoxus* windows at ≥1×, ≥2×, ≥5×, ≥10×, and ≥20×, respectively. (Fig. S1) The resulting pattern is indistinguishable in form from the contiguous tracts that an introgression caller is designed to report. (Fig. 1B)

### Mechanism 1: cytoplasmic reads with no legitimate target

To further investigate this, we rebuilt the reference adding only the *S. cerevisiae* mitochondrial genome (mtDNA; NC_001224.1) and 2-micron plasmid (NC_001398.1) – referred to as REF-cyto – and remapped the identical reads. The cytoplasmic content of the library was substantial. Under REF-cyto, 12.81 M reads mapped to the mitochondrion and 0.82 M to the 2-micron plasmid, and the overall mapping rate rose from 83.6 to 98.3%. (Fig. S2) The majority of those reads failed to map under REF-naive rather than mismapping. Approximately 0.88 M reads moved off nuclear sequence when 2-micron and mtDNA were supplied, of which ∼0.48 M had been assigned to a non-cerevisiae species. The cytoplasmic mechanism is real but smaller than the naive hypothesis predicts, because most homeless reads are lost rather than misassigned.

Its effect is nonetheless specific and directional. At 147×, adding 2-micron and mtDNA reduced the total non-cerevisiae fraction from 2.02 to 1.62%, but the reduction concentrated in the more divergent donor. *S. eubayanus*-assigned reads fell by 89% at read level (243,857 _→_ 25,850) and its window fraction by 40% (0.406% _→_ 0.245%), whereas *S. paradoxus* fell by 33% at read level (Fig. 2B) and 15% by window fraction (1.612% _→_ 1.374%). Cytoplasmic cross-mapping preferentially inflated the most divergent species in the reference, plausibly because a read with no true target is assigned by weak, near-random similarity, which is as likely to favor a distant congener as a near one.

**Figure 2.**
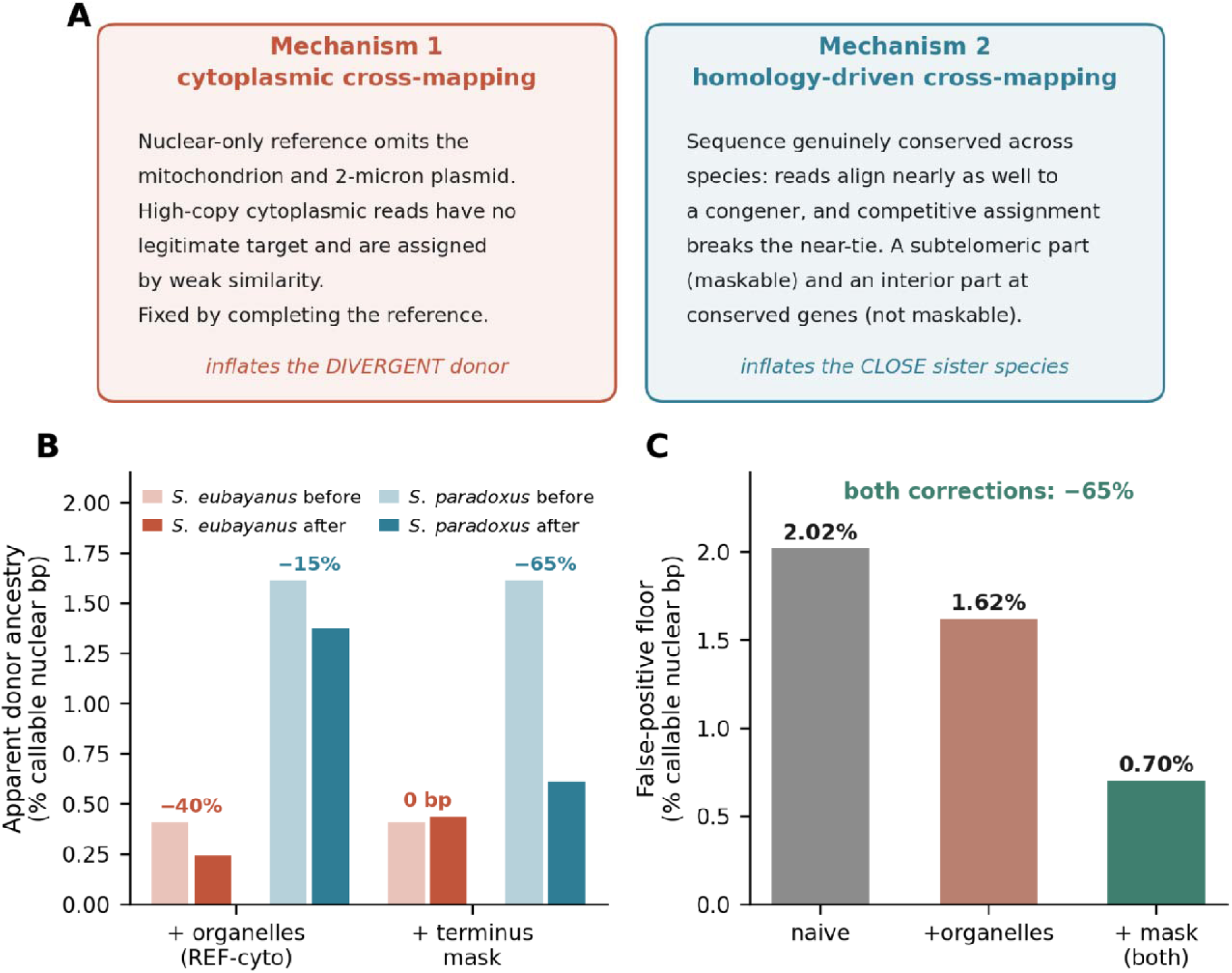
Two mechanisms with opposite phylogenetic-distance signatures. **(A)** Schematic of the two failure modes: cytoplasmic cross-mapping (reads with no legitimate target, inflating the divergent donor, fixed by completing the reference) and homology-driven cross-mapping (genuine cross-species conservation inflating the close sister, with a maskable subtelomeric component and an interior component at conserved genes that masking cannot reach). **(B)** Effect of each correction on each donor at 147×, in bp: adding *S. cerevisiae* organelles removes *S. eubayanus* signal (−40% by window fraction, −89% by read count) but little *S. paradoxus*; terminus masking removes *S. paradoxus* (−64.8% by bp) but exactly 0 bp of *S. eubayanus*. **(C)** The two corrections are additive: 2.02% _→_ 0.70%.

### Mechanism 2: homology-driven cross-mapping, only partly maskable

The residual floor under REF-cyto (1.62%) remained larger than most published introgression estimates for individual wild isolates (Barbosa, et al. 2016; Peter, et al. 2018). The surviving windows concentrated near contig ends, so we applied a terminus mask excluding the first and last 20 kb of every nuclear chromosome of every species from both numerator and denominator – a reclassification of existing coverage, with no remapping.

Masking removed more of the floor than the 2-micron and mtDNA fix had (2.02% _→_ 1.05% under REF-naive), and its species signature is the mirror image of Mechanism 1. *S. paradoxus* fell 62% (1.612% _→_ 0.610%), while *S. eubayanus* was unchanged (marginally higher: 0.406% _→_ 0.436%, an artifact of the shrinking denominator). Applied together, the two corrections reduced the floor from 2.02 to 0.70% (Fig. 2B and C; Table 1).

**Table 1.** Decomposition of the false-positive floor in a strain with no introgression. Apparent non-*S. cerevisiae* ancestry recovered from *S. cerevisiae* S288C reads (SRR2070491) under a 2×2 design: reference completeness (nuclear-only *vs*. organelle-inclusive) × terminus masking (unmasked *vs*. first/last 20 kb excluded). Values are the percent of callable nuclear bp, given in total and separately for *S. eubayanus* and *S. paradoxus*, at both the 147× and 393× conditions. The two corrections are independent and additive (2.02% _→_ 0.70% at 147×) and have opposite donor-species signatures.

| Depth | Reference | Terminus<br>mask | Total<br>non-*cerevisiae* | *Saccharomyces*<br>*eubayanus* | *Saccharomyces*<br>*paradoxus* | Callable<br>denominator (bp) |
| --- | --- | --- | --- | --- | --- | --- |
| 147× | REF-naive | unmasked | 2.0177% | 0.4059% | 1.6119% | 12,319,684 |
| 147× | REF-naive | masked | 1.0453% | 0.4355% | 0.6098% | 11,480,000 |
| 147× | REF-cyto | unmasked | 1.6184% | 0.2445% | 1.3739% | 12,269,684 |
| 147× | REF-cyto | masked | 0.6993% | 0.2622% | 0.4371% | 11,440,000 |
| 393× | REF-naive | unmasked | 3.8534% | 0.7169% | 3.1365% | 12,554,892 |
| 393× | REF-naive | masked | 2.6564% | 0.7712% | 1.8852% | 11,670,000 |
| 393× | REF-cyto | unmasked | 3.0035% | 0.2411% | 2.7625% | 12,444,892 |
| 393× | REF-cyto | masked | 1.8998% | 0.2591% | 1.6408% | 11,580,000 |

Masking removes 64.8% of *S. paradoxus* bp under REF-naive and 70.3% under REF-cyto, while *S. eubayanus* is unchanged in bp (its fraction rises only because the callable denominator shrinks). Organelle completion removes 40.0% of *S. eubayanus* bp but only 15.1% of *S. paradoxus* bp. The two corrections target different mechanisms and are additive.

The two mechanisms are independent, additive, and diagnosable by which donor they inflate. 1) Cytoplasmic cross-mapping inflates divergent donors, is fixed by reference completion, and is unaffected by masking. 2) Homology-driven cross-mapping inflates closely related donors, is partially fixed by masking, and is unaffected by reference completion. (Fig. 2A) The basis of the second mechanism is genuine cross-species sequence conservation. Where *S. cerevisiae* and a congener are nearly identical, reads align almost equally well to both, and competitive assignment breaks the near-tie without evidence. Such conservation is enriched at subtelomeres, which is why terminus masking works as well as it does, but it is not confined to them, and that distinction turns out to matter.

A replicate down-sampling experiment localized the maskable, subtelomeric part of this mechanism precisely. Repeating the 147× down-sample under five independent random seeds produced only two distinct outcomes (2.0177% and 2.0591%), differing by a single ∼5.2 kb terminal partial window of *S. paradoxus* crossing the calling threshold (+5,208 bp *S. paradoxus*, 0 bp *S. eubayanus*, +5,208 bp callable denominator; Fig. S3). Under terminus masking, all seeds returned bit-for-bit identical values (1.0453%; zero variance across six seeds, both references). The stochastic component of the floor therefore lives entirely in threshold-marginal subtelomeric windows.

### A second, more divergent control

To test whether these findings generalize beyond the reference strain itself, and to probe the interior residual where masking cannot reach, we added a second control: EM14S01-3B, a wild *S. cerevisiae* isolate from a phylogenetically divergent Asian lineage (Peter, et al. 2018), sequenced to ∼267× and roughly 1% divergent from the S288C reference (2.4× the residual divergence of S288C against its own reference). This strain plays two roles, which are worth noting here. 1) For *S. eubayanus* – an ancestry no wild *S. cerevisiae* has a plausible route to acquire – it is a second negative control, and it behaves like one. After both corrections, its apparent *S. eubayanus* signal is 0.156%, below S288C’s 0.245%. 2) For *S. paradoxus* it is not a clean negative control because, like many wild *S. cerevisiae*, it carries genuine *S. paradoxus* introgression (Barbosa, et al. 2016; Peter, et al. 2018). That introgression is what we exploit below, where its confirmed-artifact windows serve as the specificity test set for the locus-level controls (sensitivity is measured separately, on an independent strain with published, phylogenetically confirmed introgression). Its larger, more divergent *S. paradoxus* signal also makes it the better strain with which to characterize the interior, non-maskable component of the homology mechanism, because S288C’s own interior residual is only two windows.

### Masking has an optimum and cannot reach zero

If the homology mechanism were purely subtelomeric, widening the terminus mask would eventually drive the artifact to zero. It does not. Sweeping the mask from 20 to 200 kb per chromosome end, the residual *S. paradoxus* signal falls and then plateaus at a non-zero floor: 20,000 bp (2 windows) for S288C from a mask width of 100 kb onward, and 120,000 bp (12 windows) for EM14S01-3B from 150 kb onward. Neither reaches zero at 10× the standard mask width. The surviving windows are the blocks sitting 110–510 kb from any chromosome end, unreachable by any subtelomeric definition (Fig. 3A).

**Figure 3.**
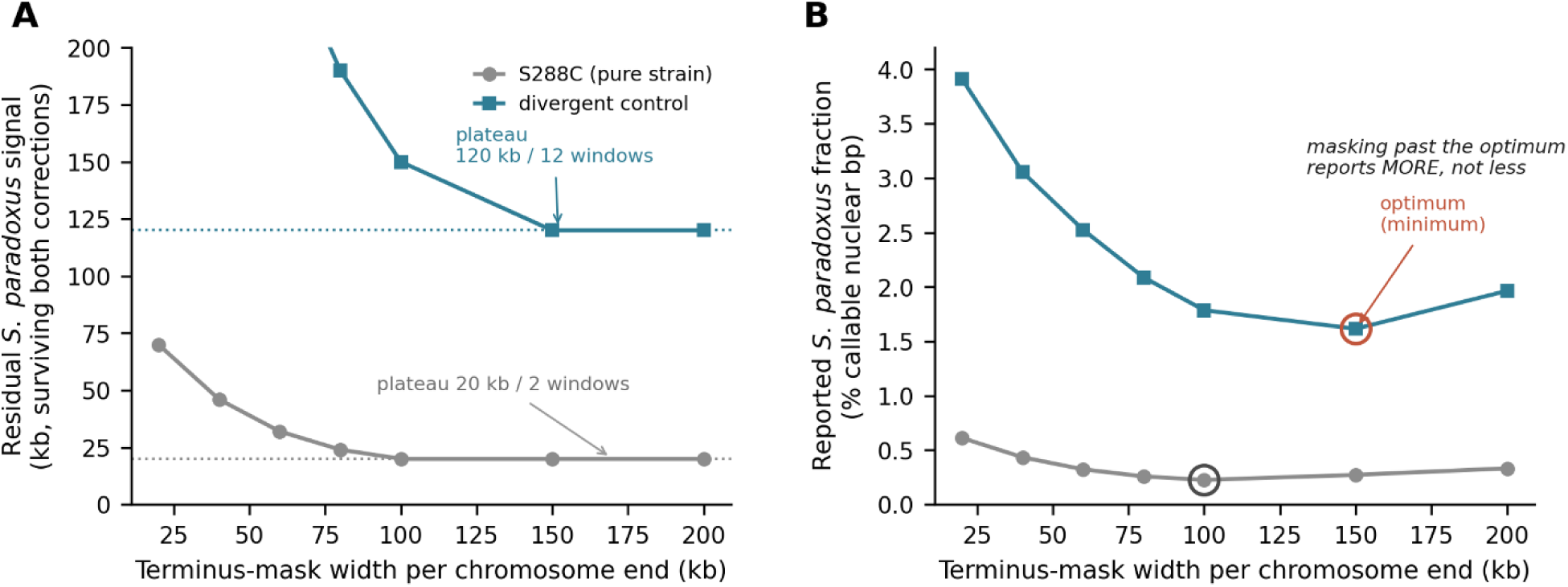
Terminus masking has an optimum and cannot reach zero. **(A)** Residual *S. paradoxus* signal surviving both corrections *vs*. terminus-mask width (20–200 kb per chromosome end), for the pure-strain control (S288C) and a more divergent control. Both fall to a non-zero plateau – 20,000 bp / 2 windows for S288C from 100 kb, 120,000 bp / 12 windows for the divergent control from 150 kb – and neither reaches zero at 10 times the standard 20 kb mask. **(B)** The same data as the reported percentage: because the callable denominator shrinks faster than the residual, the fraction reaches a minimum at the plateau onset and then rises, so masking beyond the optimum reports more apparent introgression while discarding up to half the genome. Some intermediate mask-width points are interpolated between measured endpoints.

Worse, expressed as the reported fraction, masking beyond the plateau is actively counterproductive. Because the callable denominator keeps shrinking while the residual numerator is flat, the *S. paradoxus* fraction reaches a minimum and then rises. For S288C, this is 0.437% at the standard 20 kb mask, a minimum near the plateau onset, then back up to 0.332% at 200 kb, by which point 50.9% of the callable genome has been discarded (Fig. 3B). There is an optimal mask width, the plateau onset, beyond which one loses genome and reports more apparent introgression, not less. Terminus masking is, therefore, a useful but bounded positional proxy for the underlying problem, not a solution to it. It works partly because conserved sequence clusters subtelomerically, and it fails in the interior no matter how far it is extended.

### The interior residual is a conserved-locus mapping artifact, not introgression

To identify the interior residual directly, using EM14S01-3B where it is largest, we assembled the reads underlying it and placed each locus phylogenetically. The affected loci are not repeats: of 19 assembled loci, most are ordinary single-copy genes at genome-median depth, and a per-locus test resolves their origin unambiguously. Extracting each locus’s true sequence and building a tree with the S288C and *S. paradoxus* CBS432 orthologs plus a *S. eubayanus* outgroup, 11 of 12 tested loci group with *S. cerevisiae*. The control’s own allele is *cerevisiae*-type, and the apparent *paradoxus* signal is a mapping artifact, not retained ancestral polymorphism or introgression. The twelfth, spanning *YDR035–037W*, returned an anomalous topology at low bilateral alignment coverage and coincides with a documented misassembly on chromosome IV of the CBS432 reference (Yue, et al. 2017); we exclude it as uninterpretable rather than assign it.

What these loci share is unusually high *S. cerevisiae*-*S. paradoxus* identity for their position in the genome. Two of them are *ENO2* and *TEF2*, retained duplicates from the whole-genome duplication (Wolfe and Shields 1997; Kellis, et al. 2004) and among the most highly expressed genes in the genome (Holstege, et al. 1998; Nagalakshmi, et al. 2008). They sit above the 99th percentile of genome-wide cross-species identity, and several more are moderately elevated. At such loci, a short read carries, on average, well under one base of information distinguishing the two species (the per-locus identity margins run from 0.29 to a few percentage points, *i.e*., 0.3 to a few discriminating bases per 100-bp read), so competitive assignment is close to a coin toss and resolves by read-sampling noise. This is the same homology mechanism as the subtelomeric component genuine cross-species similarity that competitive mapping cannot resolve, appearing at conserved interior loci where masking cannot reach it.

### Two standard validation heuristics fail

Two heuristics are commonly invoked to argue that a candidate introgression call is genuine. Neither survives contact with a negative control:

#### Long reads do not escape the artifact

We repeated the analysis using Oxford Nanopore whole-genome data from wild *S. cerevisiae* isolates with no plausible *S. eubayanus* contact, mapped with minimap2 against a cytoplasm-aware two-species reference. The floor persisted: 0.479% (73.97× median nuclear depth) and 0.643% (168.79×), again as discrete blocks (five and seven, respectively). Terminus masking removed 81.9% and 73.0% of that signal, as large as or larger than the ∼62% removed in short-read data (Fig. 4A). On reflection, this is expected rather than surprising. Read length rescues mapping when ambiguity arises from local repeat structure a longer read can span. It does not help when the ambiguity arises because the sequence is genuinely similar between species, which is the situation at *Saccharomyces* subtelomeres. Long reads mitigate repeat-driven mismapping; they do not mitigate homology-driven mismapping.

**Figure 4.**
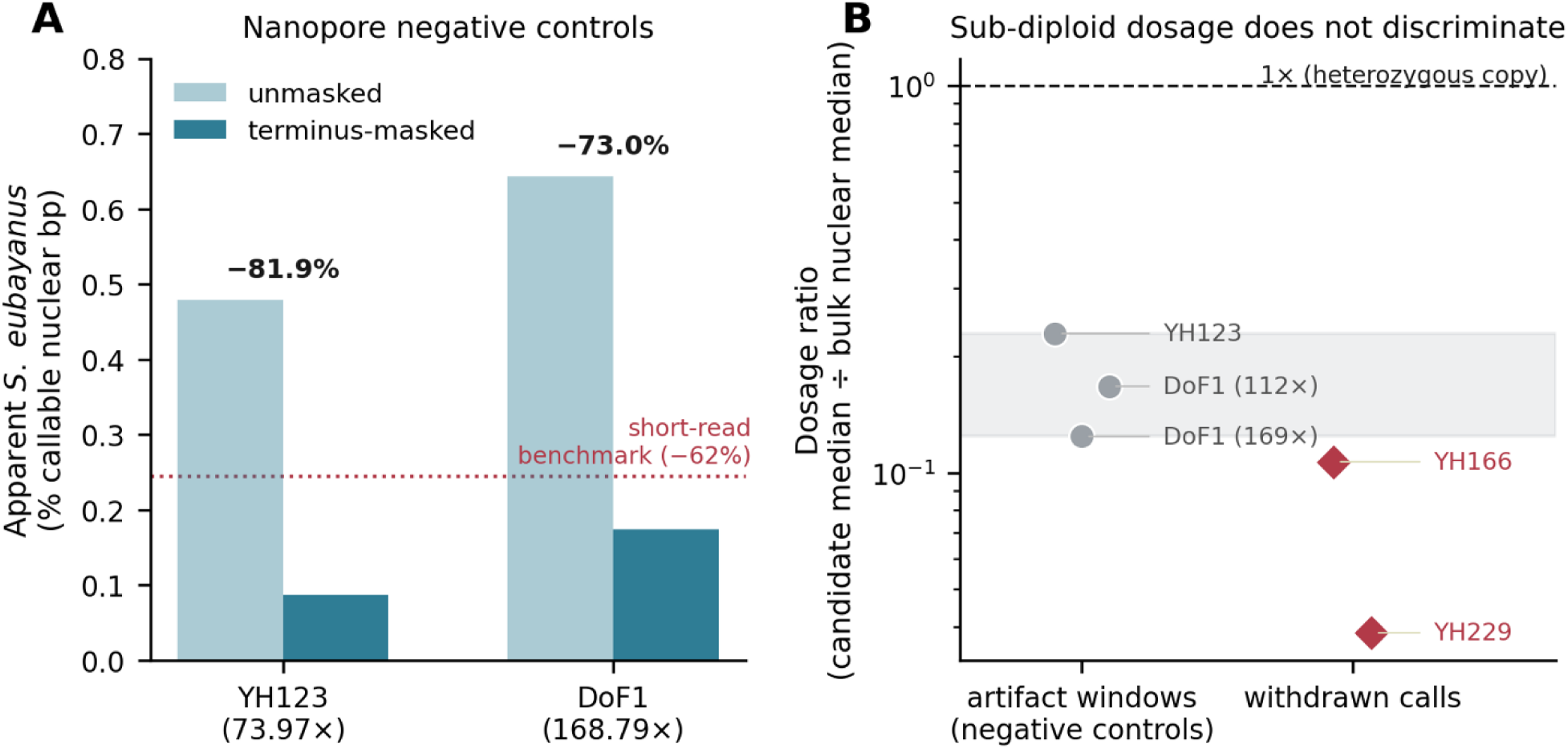
Long reads do not escape the artifact, and dosage does not discriminate it. **(A)** Apparent *S. eubayanus* fraction in two Nanopore negative controls, unmasked *vs*. terminus-masked; the relative reduction (81.9%, 73.0%) equals or exceeds that in short-read data (∼62%). **(B)** Dosage ratio (median depth of candidate windows ÷ bulk nuclear median) for negative-control artifact windows (0.229, 0.124) plotted against the values obtained for the two withdrawn calls (0.107, 0.039). Artifact windows are sub-diploid, so sub-diploid dosage does not distinguish artifact from introgression.

#### Sub-diploid dosage does not discriminate

Genuine introgressed material is often expected at reduced relative dosage (Barbosa, et al. 2016; Peter, et al. 2018) – in a subclonal population, or when carried on one of two homologs in a heterozygous diploid – and observing sub-diploid coverage in candidate blocks is treated as support. We computed, for each negative control, the median depth of its artifactual windows relative to its own bulk nuclear median. Artifact windows are strongly sub-diploid: dosage ratios of 0.229 and 0.124, roughly 4-fold and 8-fold below bulk, squarely within the range routinely read as evidence for real subclonal introgression (Fig. 4B). The reason the heuristic fails is that two distinct causes produce the same signature. Artifactual windows are depleted mechanically, because regions of low mapping uniqueness recruit fewer confidently assigned reads. Genuine heterozygous introgression is depleted biologically, because only one homolog carries the donor allele, and it sits near half-dosage. We confirm the latter directly below, in a strain whose introgression is heterozygous and reads at ∼0.5×. Sub-diploid dosage is therefore not evidence of authenticity. It is equally a signature of the artifact, and the two are not separable by dosage alone.

### FIDDL: comparative controls that discriminate

The controls that work share a common logic: rather than asking whether a candidate block looks like introgression, ask whether it behaves differently from what the same pipeline recovers in a genome that cannot contain the donor ancestry. We implement four such controls in FIDDL (*False Introgression Detection via Depth-matched controls and Loci-recurrence*), an open-source Python package with a command-line interface, released as v0.1.0.

*1) Depth-matched negative-control floor (FIDDL floor).* A strain that cannot contain the donor ancestry, sequenced on the same platform and chemistry, is processed through the identical pipeline. The candidate and control are down-sampled to common depths across multiple random seeds, and the apparent donor fraction, block count, and dosage ratio are reported for each, masked and unmasked. Because the floor rises with depth, comparing a candidate against an unmatched control is uninformative. As we demonstrate below, a call can appear to clear a floor measured at lower depth and fail once the comparison is matched.
*2) Exact-coordinate loci recurrence (FIDDL recurrence).* Genuine introgression is lineage-specific. Artifacts recur at the same coordinates across unrelated strains, because they are a property of the reference rather than of the sample. FIDDL reports, per candidate block, whether its exact coordinates are also called in the control set at matched depth.
*3) Per-read composition and identity (FIDDL identity).* For each candidate block, FIDDL reports whether coverage derives from primary or supplementary alignments and whether per-read alignment identity is higher against the focal or the donor genome. An alignment-score margin computed on reads already selected by competitive mapping is circular – those reads were chosen because the donor was their best-scoring target – so the score margin is retained for completeness but is not the decisive signal.
*4) Assembly representation (FIDDL assembly).* Sequence claimed to be present in a genome should be recoverable from a *de novo* assembly of the same reads. FIDDL aligns the assembly to the combined reference and reports whether candidate loci are represented.

FIDDL screen runs the four in sequence and emits a per-block keep/drop verdict. The package wraps the analysis logic used throughout this study rather than reimplementing it, reproduces the published control values to a sixth-decimal rounding artifact, and includes a bundled example that runs end to end in minutes and self-checks against committed expected output. The four controls are comparative: each asks whether a candidate resembles a matched negative. A natural question is whether the artifact can instead be removed directly, by filtering the offending reads or windows before any call is made. We tried, and it cannot, a result worth reporting because the reason it fails defines what a working control must do (see below).

### Why read-level correction fails

The depth-matched floor establishes that competitive mapping assigns donor ancestry to sequence that has none. If those misassigned reads could be identified and discarded, the artifact would be removed at the source. The interior-mechanism analysis suggests exactly how to try: the discriminating information at an affected locus is a per-read identity margin below one base. Thus, a filter that keeps only reads whose margin exceeds what the read length can resolve should discard precisely the uninformative reads. For 100-bp reads, that resolution limit is one base in 100, ∼0.98%.

Applied to the pure-strain control, this removes the entire residual, including the masking-irreducible plateau: 50,000 bp / 5 windows and the 20,000 bp / 2 window plateau both fall to zero, because 97% of the reads at those windows are individually uninformative. However, applied to EM14S01-3B, it removes only about half (390,000 bp / 39 windows _→_ 190,000 bp / 19 windows), and phylogenetic placement of the survivors shows they are still artifact: all 29 tested windows (the 19 that survived the filter and 10 that it removed) group with *S. cerevisiae*. The filter did not separate biology from artifact; it split one artifact population in two. Tellingly, the surviving windows carried smaller consensus margins than the removed ones (4.35 *vs*. 5.98 percentage points), the opposite of what a working discriminator would produce.

Two further attempts confirmed the limit rather than escaping it. Adding an absolute-identity floor to the margin criterion failed at the first check: the reads in surviving and removed windows are drawn from the same identity distribution (medians identical to three decimal places), so no threshold separates them. Building a consensus from the donor-assigned reads and placing *that* phylogenetically failed differently and more instructively. It called 50% of known-artifact windows introgressed because selecting reads by the assignment one is trying to test is circular: the consensus inherits the donor-ward bias of the reads that built it.

The common lesson is that no read-level or window-level filter can separate this artifact from genuine introgression because, at the loci that matter, the discriminating information does not exist within a single read. A working control must aggregate all reads at a locus into a single test and ask a question competitive mapping never asks, not “which species does this read resemble?” but “which species does this locus’s consensus sequence belong to?”

### Two validated locus-level controls

Aggregating over reads restores the missing information, and two such tests validate cleanly on ground truth: the 29 phylogenetically confirmed artifact windows above (specificity); and an independent set of published, phylogenetically confirmed introgression loci from a wild isolate (sensitivity).

#### Consensus-phylogenetic placement

For each candidate block, build a consensus from all reads at the locus regardless of species assignment, and place it in a tree with the focal and donor references and an outgroup. Accept the block only if the consensus groups with the donor. Because it uses every read at the locus, it is not subject to the selection bias that defeated the donor-assigned consensus. It rejected all 29 confirmed-artifact windows (29/29 specificity). Its one blind spot is the homozygous case. At a locus introgressed on both homologs, it returns a single donor-type consensus correctly, but at a *heterozygous* locus it returns the majority allele, which may be either.

#### Allele-fraction donor-match

For each candidate block, using all reads mapped to the focal reference, call heterozygous sites and ask what fraction of their alternate alleles match the donor reference base at the syntenic position. Genuine heterozygous introgression produces many heterozygous sites whose alternates systematically match the donor; artifact produces few, with no systematic donor match. The two known sets separate with an empty gap between donor-match fractions of 0.333 and 0.778. At the gap midpoint (0.5, unfitted), the test rejected all 29 artifact windows and accepted 30 of 42 resolvable introgression loci. The 12 misses are diagnostic rather than random: four are homozygous tracts (no heterozygosity to measure, the consensus test’s blind spot in mirror image), six are technical liftover failures, and two are genuine low-donor-match loci.

The two controls are therefore complementary. The allele-fraction test catches heterozygous introgression that the consensus test reports only as a majority allele, and the consensus test catches homozygous introgression to which the allele-fraction test is blind. Together, they constitute a one-sided but reliable acceptance test: a high donor-match fraction or a donor-grouping consensus is strong evidence of genuine introgression, while a negative result does not by itself exclude it. For a control whose purpose is to prevent false positives, one-sidedness in this direction is exactly what is required. Both are prototyped against the analyses reported here and are candidate additions to a future FIDDL release; their behavior at low sequencing depth, where heterozygous-site calling degrades, remains to be characterized.

### Validation: two of our own introgression calls withdrawn

We applied FIDDL to two isolates from our own collection. YH166 and YH229 are wild *S. cerevisiae* recovered from spontaneously fermented beer (companion paper, Shumaker *et al*.). Both produced reduced mapping to the S288C nuclear reference (60.5% and 77.6% primary mapping, *vs.* 93–96% for other *S. cerevisiae* isolates in the same collection), and analysis had already passed through two rounds of correction. In the first, competitive mapping against a nuclear-only combined reference indicated *S. cerevisiae* × *S. eubayanus* hybridity. In the second, supplying the *S. cerevisiae* mtDNA and 2-micron plasmid removed most of that signal and reduced the interpretation to minor, patchy nuclear introgression: 1.04% of callable nuclear positions in YH229 (10 blocks, ∼127 kb) and 0.48% in YH166 (5 blocks, ∼58 kb), at sub-diploid dosage ratios of 0.039 and 0.107. That interpretation was supported by three conventional lines of evidence – signal above background, sub-diploid dosage, and discrete block structure – and was manuscript-ready. However, it does not survive the four controls (Fig. 5).

**Figure 5.**
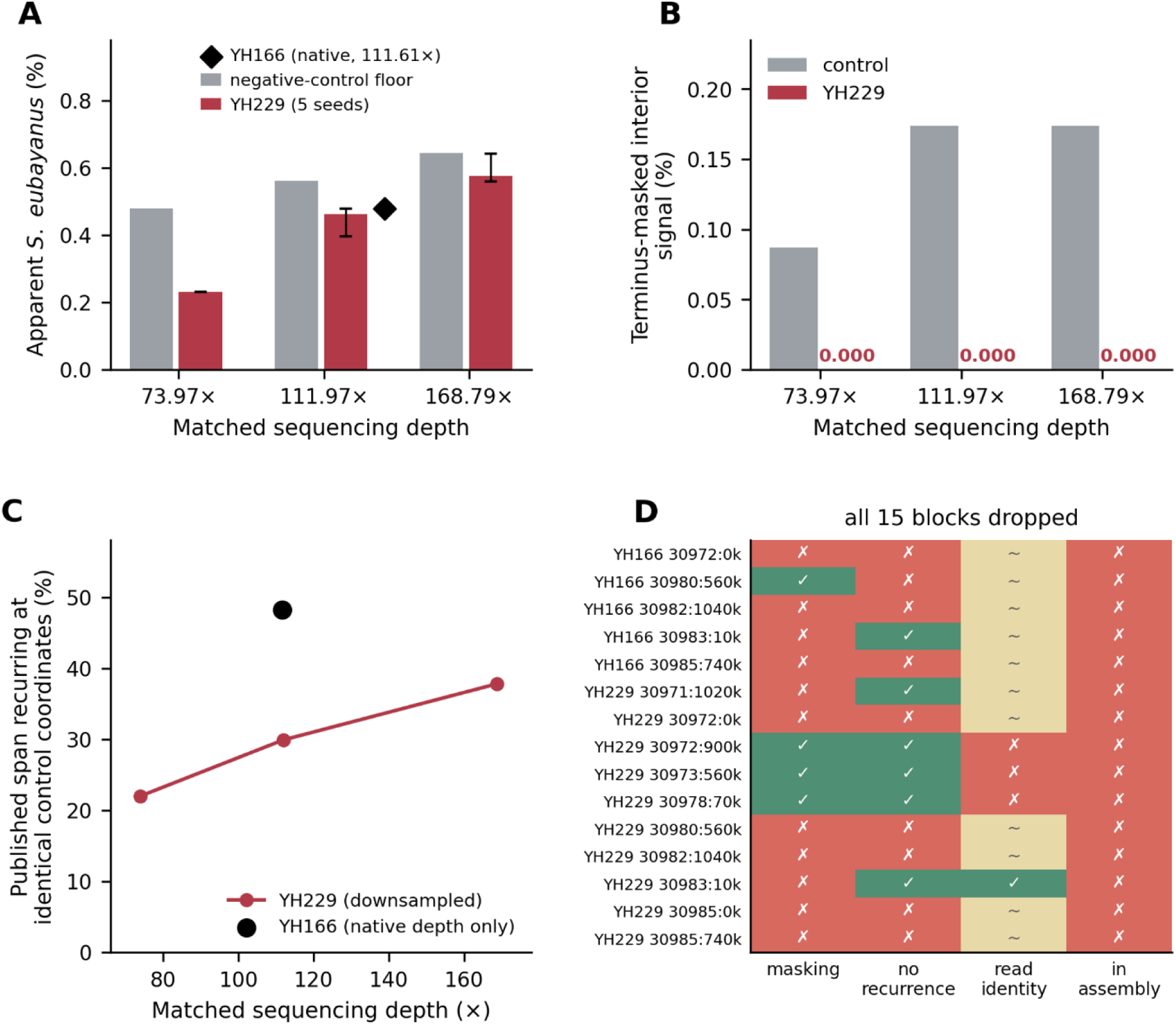
Two introgression calls withdrawn under matched controls. **(A)** Apparent *S. eubayanus* fraction for isolates and negative controls at three matched sequencing depths (YH229 down-sampled, five seeds each; mean and range). YH166 is shown at its single native depth (111.61×), as it was the depth to which the controls were matched and so was not itself down-sampled. At every matched depth, the isolates fall at or below the control floor. **(B)** Terminus-masked interior signal: zero for YH229 at all matched depths, while controls retain more. **(C)** Fraction of each isolate’s published span recurring at identical coordinates in unrelated controls *vs*. matched sequencing depth. YH229’s recurrence falls at lower depth only because fewer of its blocks remain callable there; at the lowest depth (73.97×), all of its still-callable blocks recur in the controls. YH166 is a single native-depth measurement (48.2%). **(D)** Per-block outcome against the four criteria, showing zero surviving blocks in either isolate.

#### Matched-depth floor

Down-sampling both isolates and controls to common depths (five seeds per condition) placed the isolates at or below the control floor at every depth tested: YH229 gave 0.576% at 168.79× against a floor of 0.643%, 0.461% at 111.97× against 0.561%, and 0.231% at 73.97× against 0.479%. YH166, at its native 111.61×, gave 0.479% against the same 0.561% floor. The apparently favorable 1.62× margin YH229 had shown against a lower-depth control was entirely a depth confound. (Fig. 5A)

#### Recurrence

Between 22 and 38% of YH229’s published span, and 48% of YH166’s, fell at exactly the same 10 kb coordinates called in unrelated control strains. At the lowest matched depth, all of YH229’s still-callable blocks recurred in controls. (Fig. 5C)

#### Interior signal

Terminus masking reduced YH166’s native-depth signal by 81.8%, indistinguishable from the 81.9% reduction in a negative control. YH229’s masked interior signal fell to exactly zero at every matched depth across all five seeds, while the negative controls retained more (0.174% and 0.087%). The interior residual YH229 showed at its native 420.84× exists only at that depth. (Fig. 5B)

#### Read composition and identity

Of the three YH229 blocks surviving masking and recurrence filtering, two contained zero primary reads – their entire coverage derived from supplementary alignments of reads whose primary alignments lay on S. cerevisiae chromosomes XIV and XVI. The third aligned at higher mean identity to *S. cerevisiae* (94.0%) than to *S. eubayanus* (83.6%). (Table S1)

#### Assembly

No contig of either de novo assembly maps to *S. eubayanus* anywhere in the genome, despite both assemblies being near expected size and contiguity (YH166: 12.40 Mb, 41 contigs, N50 801 kb, of which 38 map to *S. cerevisiae* nuclear and 3 to cytoplasmic sequence; YH229: 12.25 Mb, 31 contigs, N50 822 kb, 20 nuclear and 11 cytoplasmic). (Fig. S4)

No block in either isolate satisfies all four criteria; the defensible residual is 0 bp for both. One locus (NC_030983.1:10,000–20,000) fails only the positional and assembly criteria and is the single candidate the data cannot fully exclude. It lies within the terminus zone where the dominant mechanism operates (Fig. 5D). Another detail worth noting is that at 111.97×, YH229 returns 0.461%, and YH166 at 111.61× returns 0.479%. Two different libraries from two different isolates converge on the same value at the same depth, because they are measuring the same artifact rather than two independent biological findings.

### Comparison with current practice: magnitude in published data

We asked how much of the introgression reported in a recent published panel is attributable to these mechanisms. We selected sixteen wild *S. cerevisiae* isolates from (Avelar-Rivas, et al. 2026), spanning their reported range (9 to 364 blocks), remapped them under the study’s own 10-species concatenated reference design, and remapped again with both corrections applied (Fig. 6).

**Figure 6.**
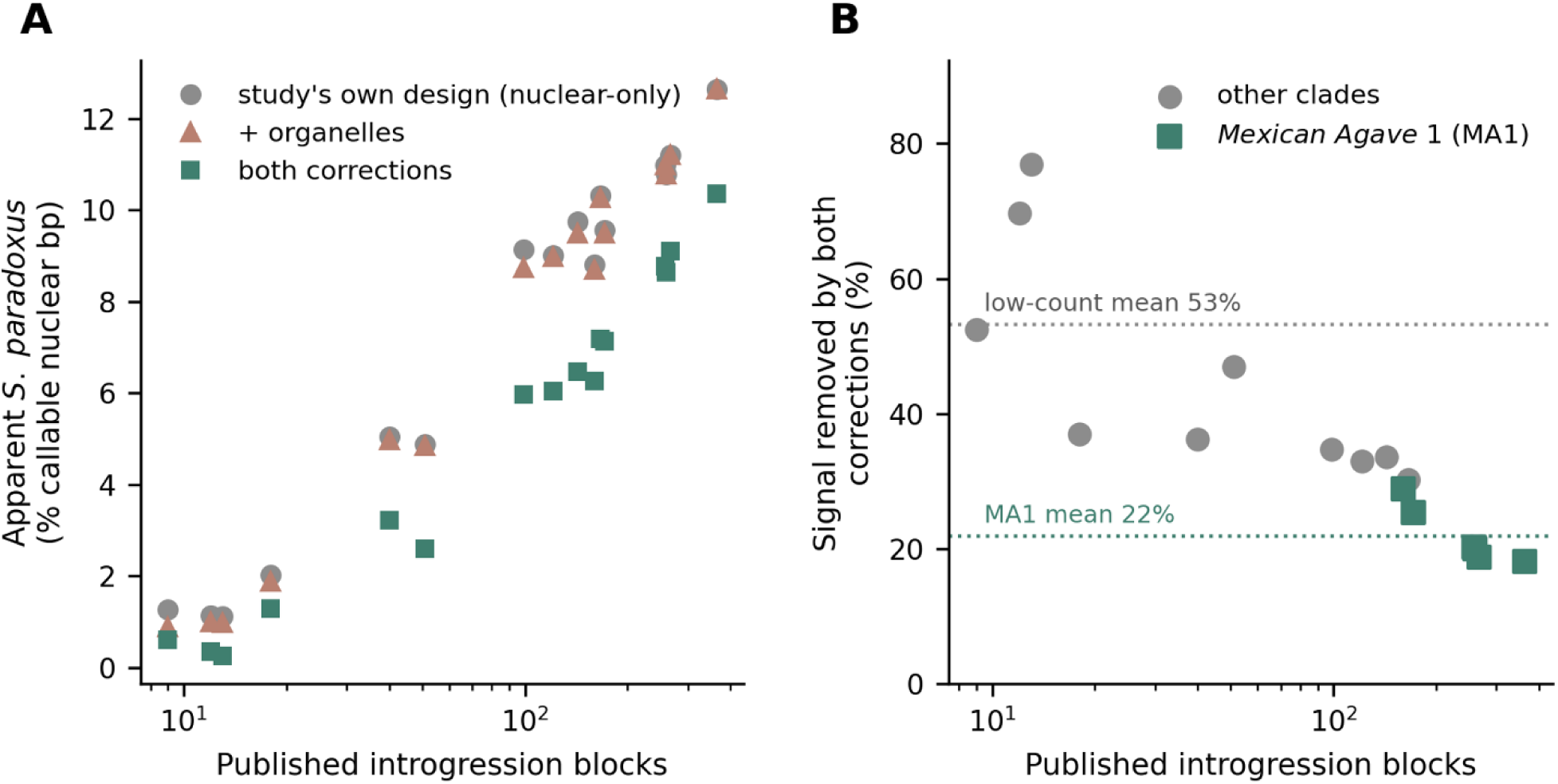
Correction magnitude in published data. For 16 wild isolates from (Avelar-Rivas, et al. 2026), apparent *S. paradoxus* fraction under the study’s own reference design, with the organelle correction alone, and with both corrections, plotted against published introgression block count. Low-count calls lose a mean of 53.2% of apparent signal; high-count MA1 calls lose 18.2-28.9% (mean 21.9%) and retain double-digit signal.

The corrections behaved as the mechanism analysis predicts. In the lowest-count isolate (YMX002071, nine reported introgressions) (Avelar-Rivas, et al. 2026), apparent *S. paradoxus* signal fell from 1.27% to 0.89% with the 2-micron/mtDNA fix alone and to 0.60% with both, a 52% reduction. Across the six lowest-count isolates (9-51 blocks) the mean reduction was 53.2% (range 36.2-76.9%). In the six isolates of the Mexican Agave 1 (MA1) clade, which carry the study’s highest counts (160–364 blocks), the 2-micron/mtDNA fix alone was negligible (0.0-1.2%), and both corrections together removed 18.2–28.9% (mean 21.9%), leaving double-digit apparent introgression intact. The highest-count isolate fell from 12.65 to 10.35%.

We also tested whether the terminus-proximal component in these isolates might be genuine subtelomeric introgression rather than artifact. Real introgression should recur at shared coordinates within related strains and be absent from unrelated ones (Peter, et al. 2018). The near-terminus signal instead appears in 100% of MA1 strains and in unrelated low-introgression clades – a clade-agnostic distribution characteristic of artifact rather than lineage-specific biology.

These results are not a claim that the published counts are wrong. That study calls introgression at gene level and requires a divergence threshold and phylogenetic confirmation against real *S. paradoxus* references before reporting a block (Avelar-Rivas, et al. 2026). That is a materially stronger filter than the coverage-threshold approach used here, and one that artifactual sequence, having no true donor ancestry, would be expected to fail. What the re-analysis quantifies is the magnitude of the vulnerability in the general first-pass approach and, equally, that this study’s phylogenetic safeguard is doing necessary work. The pattern that matters is the contrast: low-confidence, diffuse calls are substantially inflated; large, gene-level, phylogenetically confirmed tracts are robust.

## DISCUSSION

### What is and is not at risk

The results reported here should not be read as an indictment of introgression genomics. Large, contiguous, gene-level introgression tracts – the Alpechin *S. paradoxus* introgression (D’Angiolo, et al. 2020), lager-type hybrid subgenomes (Dunn and Sherlock 2008; Libkind, et al. 2011), and the high-count MA1 signal (Avelar-Rivas, et al. 2026) examined here – are supported by coverage far above any floor we measured, survive both corrections, and are typically confirmed phylogenetically. Nothing here refutes them.

What is at risk is the tail: low-level, spatially diffuse, subtelomerically distributed calls reported without a matched negative control, especially from deeply sequenced libraries. That tail is not a negligible part of the literature (Peter, et al. 2018; Avelar-Rivas, et al. 2026). It is where claims of “residual,” “minor,” or “cryptic” introgression live, and where the interesting evolutionary interpretations – ancient contact, adaptive retention, permeable species boundaries – are typically drawn. Our own withdrawn calls sat precisely there.

### Three assumptions that do not hold

Each principal finding contradicts a working assumption, including assumptions we held ourselves while generating the calls we later withdrew. 1) More depth is not more reliable. The floor was negligible at 10-25× and rose continuously through 393×. A deeply sequenced genome is more, not less, likely to produce spurious interspecific blocks, and the effect persists under a depth-scaled threshold. Studies that sequence a subset of strains more deeply, or compare across libraries of unequal depth, will find apparent introgression tracking sequencing effort. 2) Long reads are not a solution. The subtelomeric mechanism was at least as strong in Nanopore as in Illumina data. Read length addresses ambiguity from repeat structure, not from genuine cross-species similarity, and at *Saccharomyces* subtelomeres, the similarity is real. 3) Sub-diploid dosage is not evidence. Depleted coverage in candidate blocks is produced mechanically by low mapping uniqueness and is reproduced by strains that cannot contain donor sequence. Used as a validation heuristic, it selects for artifact as readily as for biology.

### How easily the error is committed

We can offer direct evidence for how difficult this failure mode is to avoid, because we committed it again while building the tool designed to detect it. The depth labels in our own first-pass analysis were wrong by a factor of approximately three. The function computing mean nuclear depth excluded only the two cytoplasmic contigs, leaving the *S. eubayanus* and *S. paradoxus* nuclear chromosomes in the denominator. Because those two species together comprise roughly two-thirds of the concatenated reference and carry essentially no true depth in a pure *S. cerevisiae* control, the reported depth was diluted to about one-third of the real focal-species value. A separate function used elsewhere in the same project restricted correctly to focal contigs, so the two branches were never computing the same quantity (companion paper, Shumaker *et al*.). The down-sampling target was computed against the diluted metric as well, so a condition labeled “50×” was in fact 147×. The error was internally consistent and therefore invisible: every number was reproducible, and none was right.

Independently, during development of FIDDL itself, the routine computing depth-weighted means was found to lack a species restriction, so the tool’s own down-sampling logic was briefly calibrating against an all-species-diluted metric, *i.e.*, the same error, in the same project, inside the software written to catch it. It was found not by code review but by running the bundled example on realistic data. We report this because it bears on how the recommendations below should be read. The failure mode is not exotic and is not a matter of carelessness. It is a natural consequence of computing a species-specific quantity over a multi-species reference, and it survives review by people actively looking for it. Controls that compare against an empirical negative catch it; inspection does not.

### Recommended practice

None of the following requires new sequencing. All steps are implemented in or prototyped against FIDDL:

*1) Complete the reference or mask cytoplasmic reads deliberately.* Include the focal species’ mitochondrial genome and plasmids, or mask cytoplasmic reads before mapping, and report which was done. This removes the divergent-donor mechanism.
*2) Run a depth-matched negative control.* Use a strain that cannot contain the donor ancestry, sequenced using the same platform and chemistry, down-sampled to the same depth, through the identical pipeline. Report the floor alongside the calls. This is the single most informative control and the hardest to compensate for later.
*3) Mask termini, but do not over-mask.* Terminus masking removes the subtelomeric component of the homology mechanism, but the residual plateaus at a non-zero floor. The reported fraction rises again once masking passes its optimum. Report the mask width and treat masking as a partial, bounded correction rather than a cure.
*4) Confirm survivors at the locus level, not the read level.* Read-level and window-level filters cannot separate this artifact from genuine introgression, because the discriminating information is below single-read resolution. Aggregate all reads at a candidate locus and test the consensus phylogenetically. Where the locus is heterozygous, test whether its heterozygous-site alternate alleles match the donor. A block that survives a depth-matched floor but whose consensus groups with the focal species is an artifact.

Reporting the calling threshold’s behavior across the depth range actually used, stating library cytoplasmic content, and specifying which depth metric is quoted all cost nothing and let readers assess exposure directly.

### Limitations

The short-read floor’s two mechanisms and the read-level dead-end were characterized primarily on two control strains: a zero-divergence pure strain (S288C, the reference) and one more divergent isolate. A broader panel spanning a range of divergence from the reference would more firmly establish how the interior floor scales with reference distance, which our two points can only suggest. The interior-mechanism per-locus test rests on 12 loci, of which one was excluded as uninterpretable. Extending it would strengthen the claim that the residual is uniformly a conserved-locus artifact. The two validated locus-level controls are one-sided acceptance tests: a negative result does not exclude introgression; and their behavior at low sequencing depth, where heterozygous-site calling degrades, is untested. The consensus and allele-fraction tests have complementary blind spots (homozygous and no-heterozygosity loci, respectively) that we characterize but do not fully close. Our read-fate accounting for the short-read arm is aggregated by species rather than tracked per read. The published-panel re-analysis covers 16 isolates from one study (Avelar-Rivas, et al. 2026) and uses a coverage-threshold caller rather than that study’s full gene-level pipeline, so it bounds the vulnerability of the general approach rather than reproducing their calls. Finally, our negative controls are valid for *S. eubayanus*, which wild North American oak *S. cerevisiae* has no plausible route to acquire (Libkind, et al. 2011; Peter, et al. 2018). They are not clean negative controls for *S. paradoxus*, because introgression from *S. paradoxus* has been documented in many wild *S. cerevisiae* lineages (Barbosa, et al. 2016; Peter, et al. 2018). Indeed, our divergent control carries genuine *S. paradoxus* introgression, which is why it serves here as a positive rather than a negative control for that donor.

## Conclusion

The standard first-pass method for detecting interspecific introgression assigns 2% of a pure genome’s callable positions to other species at 147× coverage, in discrete blocks, at sub-diploid dosage, more so at greater depth, and no less so with long reads. The artifact resists the obvious fixes. Completing the reference removes only the divergent-donor component. Masking removes only the subtelomeric part of the rest and becomes counterproductive if pushed. No read-level filter can reach the remainder because at the conserved loci that generate it, a single read carries less than one base of discriminating information. What works is to stop asking whether individual reads resemble the donor and instead test, locus by locus, whether the aggregated sequence belongs to it. We report two of our own calls withdrawn on this basis, not as a curiosity, but because it is the most direct evidence we can offer that the failure mode is reachable by careful work. The controls that expose it are cheap and now packaged; the inferences that rest on skipping them are not.

## MATERIALS AND METHODS

### Depth metrics

Short-read depths are bp-weighted means over focal-species nuclear 10-kb windows. Long-read depths are medians over the same windows. The two differ by 13-15% on the same data and are reported separately throughout.

### Short-read negative control

*S. cerevisiae* S288C whole-genome Illumina data (SRA run SRR2070491; BioProject PRJNA287442; HiSeq 2000, paired-end; 42,937,674 reads, 6,396,260,614 bases; metadata verified via the ENA file report API) were mapped with bwa-mem2 (default parameters) (Vasimuddin, et al. 2019), sorted and duplicate-marked with samtools, and summarized in 10 kb windows with mosdepth --by 10000 --no-per-base (Pedersen and Quinlan 2018).

Two references were built, differing only in the presence of *S. cerevisiae* organellar sequence. REF-naive comprised the 16 nuclear S288C chromosomes, the *S. eubayanus* FM1318/CBS 12357 nuclear chromosomes, and the *S. paradoxus* CBS432 nuclear chromosomes (GCF_002079055.1). REF-cyto added the *S. cerevisiae* mitochondrion (NC_001224.1) and 2- micron plasmid (NC_001398.1). Contigs were renamed <species>__<contig> at build time for unambiguous per-window assignment.

A window was called donor-derived at ≥5× mean depth. The reported fraction is callable non-*cerevisiae* bp ÷ (callable *cerevisiae* bp + callable non-*cerevisiae* bp), bp-weighted. Analyses were run at the library’s native depth (393×) and at a down-sampled condition (147×). The 50× value quoted in Results is read from the coverage ladder’s own 49.99× rung, a separate down-sample series, at seed 0.

### Down-sampling and replicate seeds

Down-sampling used samtools view -b -s <seed>.<fraction> (Danecek, et al. 2021). In samtools view -s FLOAT, the integer part is the seed, and the fractional part the proportion. Thus, an invocation with a bare fraction is implicitly seed 0. Values reported as reproducing earlier single-draw analyses use seed 0; replicate spreads use seeds 1–5, reported as mean and full range.

### Terminus masking

The terminus mask excludes the first and last 20,000 bp of every nuclear chromosome of every species from both the numerator and denominator. It is applied as a reclassification of existing windowed coverage; no remapping is performed. Masked and unmasked classifications crossed with REF-naive and REF-cyto form the 2×2 design in Table 1.

### Depth ladder and spike-in titration

For the coverage ladder, the control library was down-sampled to nominal 10×, 25×, 50×, 100×, and 200× focal-species depth and reclassified under both references: masked and unmasked. For the composition titration, organellar reads were spiked into a fixed nuclear read set at 0-40% of total reads. Results are reported on a read-count metric, with the windowed metric shown for comparison and noted as quantized because small changes in cytoplasmic content move no window across the calling threshold.

### Long-read arm

Oxford Nanopore data were generated by Plasmidsaurus Inc. (Louisville, KY, USA) by amplification-free tagmentation (kit-14/v14 chemistry, R10.4.1 flow cells) and base-called with dna_r10.4.1_e8.2_400bps_sup@v4.3.0. Negative controls were two wild *S. cerevisiae* isolates from tree bark (YH123, 73.97×; DoF1, 168.79×) with no plausible *S. eubayanus* contact (companion paper, Shumaker *et al*.); DoF1 reads were retrieved from SRA run SRR39135318. Reads were mapped with minimap2 -ax map-ont --secondary=no (Li 2018) against a cytoplasm-aware reference (S288C 16 nuclear chromosomes + NC_001224.1 + NC_001398.1 + *S. eubayanus* FM1318 16 nuclear chromosomes), summarized with mosdepth --by 10000 (Pedersen and Quinlan 2018), and classified with the ≥5× convention above.

Down-sampling reduces depth only, so no control could be matched to YH229’s native 420.84×; the isolate was instead down-sampled to each control’s depth. For the scaled-threshold test, classification was repeated with the calling threshold set to 5% of each library’s own focal-species median depth rather than a fixed 5×.

### Block-level criteria

Four criteria were applied to every candidate block: 1) survival of the terminus mask; 2) absence of the same coordinates from the union of control blocks at matched depth, across all seeds; 3) a consistent *S. eubayanus* preference on per-read realignment against single-species references; and 4) representation in the isolate’s *de novo* assembly, assessed by aligning the assembly to the combined reference with minimap2 -x asm10 (Li 2018). For criterion (3), an alignment-score margin computed on reads already selected by competitive mapping is circular; those reads were chosen because the donor was their best-scoring target. We therefore relied on two non-circular signals: whether a block’s coverage derives from primary or supplementary alignments, and whether per-read alignment identity is higher against the focal or the donor genome.

### Published-panel re-analysis

Sixteen isolates from (Avelar-Rivas, et al. 2026) (SRA runs listed in Table S1), spanning 9–364 reported introgression blocks, were remapped under a reconstruction of that study’s 10-species concatenated nuclear reference (*S. cerevisiae* S288C, *S. paradoxus* YPS138, *S. mikatae* IFO1815, *S. kudriavzevii* Cr85, *S. jurei* M1, *S. arboricola* H6, *S. uvarum* CBS7001, *S. eubayanus* FM1318, plus *Kluyveromyces marxianus* and *Pichia kudriavzevii*), and again with *S. cerevisiae* organelles added and the terminus mask applied.

### Terminus-mask width sweep

For both the pure-strain control and the divergent control, classification under REF-cyto was repeated at terminus-mask widths of 20, 40, 60, 80, 100, 150, and 200 kb per chromosome end, reclassifying existing windowed coverage with no remapping. For each width, we recorded the callable denominator, the surviving *S. paradoxus* and *S. eubayanus* bp and window counts, and the resulting fraction. The plateau was defined as the mask width beyond which surviving bp did not decrease. The fraction minimum was located separately because the fraction rises past the plateau as the denominator continues to shrink.

### Interior-mechanism per-locus test

Reads underlying the interior residual of the divergent control (blocks surviving REF-cyto with the 20 kb terminus mask) were extracted and assembled with SPAdes (Bankevich, et al. 2012). Assembled contigs were aligned to the S288C and *S. paradoxus* CBS432 references. Loci were annotated against the S288C feature set, and per-locus copy number was estimated from depth relative to the genome-wide median. For each of 12 candidate loci, all reads at the locus (from the naive-reference alignment, without selection by species assignment) were used to build a consensus, which was aligned with the S288C and CBS432 orthologs and an *S. eubayanus* outgroup ortholog. Trees were built with FastTree (Price, et al. 2010) and topology recorded, alongside per-reference alignment identity, length, and query coverage. One locus (*YDR035–037W*) returned an anomalous outgroup-grouping topology at low bilateral coverage, coincident with a reported chromosome IV misassembly in CBS432 (Yue, et al. 2017), and was excluded as uninterpretable. A genome-wide S288C *vs*. CBS432-windowed identity distribution was computed to place each locus’s cross-species identity as a percentile and to screen CBS432 for *S. cerevisiae* contamination (assessed by whether high-identity regions clustered, as contamination would, or dispersed across contigs and chromosomes, as conserved-sequence homology does).

### Read-level and locus-level discriminators

Three read-level filters were evaluated against the 29 phylogenetically confirmed artifact windows. The margin filter retained only reads whose per-read identity margin between the focal and donor references exceeded the read-length resolution limit (1/read-length). The two-parameter filter added an absolute best-identity floor derived per strain from that strain’s genome-wide read-identity distribution. The donor-assigned-consensus test built a consensus from only the donor-assigned reads at each locus and placed it phylogenetically. Two locus-level controls were then evaluated. The consensus-phylogenetic test built a consensus from all reads at the locus and accepted the block only if it grouped with the donor. The allele-fraction donor-match test called heterozygous sites among all focal-reference-mapped reads at the locus and computed the fraction of alternate alleles matching the donor base at the syntenic position. A threshold was taken at the midpoint of the empty gap between the artifact and introgression distributions, without fitting. Specificity was measured on the 29 confirmed-artifact windows. Sensitivity was measured on an independent set of published, phylogenetically confirmed *S. paradoxus* introgression loci from a wild isolate, testing the donor-assigned allele at each heterozygous locus.

### Implementation

FIDDL is a Python package with a command-line interface, comprising modules for reference construction, per-window classification, matched-depth floor estimation, loci recurrence, read composition and identity, assembly representation, and report assembly. It wraps the analysis logic used throughout this study rather than reimplementing it. Reproduction of the published short-read and long-read control values by the packaged tool agrees to a sixth-decimal rounding artifact. A bundled example dataset runs end to end and self-checks against committed expected output. The four comparative controls are released in v0.1.0; the consensus-phylogenetic and allele-fraction donor-match tests are prototyped against the analyses reported here and are candidate additions to a future release. FIDDL v0.1.0 is archived at https://doi.org/10.5281/zenodo.21509544.

### Tool versions

The following software versions were used, unless otherwise noted: bwa-mem2 2.3, minimap2 2.31-r1302, samtools 1.23.1, mosdepth 0.3.14, bedtools 2.31.1, and python 3.11.

## Data availability

FIDDL is available at https://github.com/Bochman-Yeast/FIDDL (release v0.1.0) and permanently archived at https://doi.org/10.5281/zenodo.21509544. Reference build scripts, per-window classifications, and all intermediate tables are deposited in the same archive. Public data re-analyzed here are available under the accessions cited above; long-read data for the isolates used as negative controls and as the worked example are available under [CALL-OUT: BioProject].

## Acknowledgments

We thank members of the Bochman lab for critically reading this manuscript.

## AI use disclosure

Large language model assistance (Claude, Anthropic) was used during manuscript preparation, including designing and generating figures and reviewing the internal consistency of reported values across analyses. An agentic coding assistant built on the same model family (Claude Code) was used to execute bioinformatics analyses on Indiana University’s Quartz high-performance computing infrastructure – running alignment, classification, and phylogenetic-placement scripts and exporting result tables – under task specifications written by the authors that fixed the analysis parameters and required verbatim, read-only reporting of outputs. All hypotheses, experimental designs, and interpretive conclusions were formulated by the authors; the AI coding assistant’s role was limited to executing specified analyses and reporting results. All figures, tables, and reported values were reviewed and verified by the authors against source data before inclusion. No AI tool is an author of this work, and the authors take full responsibility for its content and accuracy.

