## Supplemental materials for "FIDDL: depth-matched negative controls distinguish genuine interspecific introgression from competitive-mapping artifact"

**Supplementary Materials**

**Supplementary Tables**

**Table S1. Per-block evidence for the two withdrawn calls (YH166, YH229).**

**Table S2. Sixteen re-analyzed panel isolates.**

**Table S3. Reference genome assemblies.**

**Table S4. Negative controls and worked example.**

**Supplementary Figures and Figure Legends**

**Figure S1. Threshold sensitivity: window counts retained at ≥1×, ≥2×, ≥5×, ≥10×, ≥20× for each donor.**

**Figure S2. Read-fate accounting for the short-read control: read counts by destination under each reference.**

**Figure S3. The residual stochasticity of the floor is a single terminal window.**

**Figure S4. Assembly cross-check: contig assignment by species for both withdrawn isolates.**

**Supplementary References**

**Table S1. Per-block evidence for the two withdrawn calls (YH166, YH229).** Every candidate *S. eubayanus* block from the original YH166/YH229 calls, evaluated against the four block-level criteria. The matched-depth floor criterion is evaluated at the isolate level, not per block (Methods): YH229 fell below the DoF1 floor at all three matched depths (0.576% *vs*. 0.643% at 168.79×; 0.461% *vs*. 0.561% at 111.97×; and 0.231% *vs*. 0.479% at 73.97×), and YH166 fell below the same floor at its native depth (0.479% *vs*. 0.561% at 111.61×). No block in either isolate passes all four criteria; every block is withdrawn. The one locus present in both isolates (NC_030983.1:10,000-20,000) fails masking and assembly in both. While its YH229 copy shows the strongest read-identity margin of any block tested (2139.3), that margin metric is circular by construction (Aim 4) and is not treated as a pass on its own.

| **Isolate** | **Contig** | **Start** | **End** | **Size (bp)** | **Terminus mask^†^** | **Coordinate recurrence^‡^** | **Read composition & identity^§^** | **Assembly representation** | **Verdict** |
| --- | --- | --- | --- | --- | --- | --- | --- | --- | --- |
| YH166 | NC_030972.1 | 0 | 20,000 | 20,000 | FAIL | recurs | not evaluable (moot) | absent | *withdrawn* |
| YH166 | NC_030980.1 | 560,000 | 570,000 | 10,000 | PASS | recurs | not evaluable (moot) | absent | *withdrawn* |
| YH166 | NC_030982.1 | 1,040,000 | 1,050,000 | 10,000 | FAIL | recurs | not evaluable (moot) | absent | *withdrawn* |
| YH166 | NC_030983.1 | 10,000 | 20,000 | 10,000 | FAIL | unique | not evaluable (moot) | absent | *withdrawn* |
| YH166 | NC_030985.1 | 740,000 | 747,934 | 7,934 | FAIL | recurs | not evaluable (moot) | absent | *withdrawn* |
| YH229 | NC_030971.1 | 1,020,000 | 1,030,000 | 10,000 | FAIL | unique | non-decisive (margin 57.4) | absent | *withdrawn* |
| YH229 | NC_030972.1 | 0 | 20,000 | 20,000 | FAIL | recurs | non-decisive (ambiguous) | absent | *withdrawn* |
| YH229 | NC_030972.1 | 900,000 | 910,000 | 10,000 | PASS | unique | fail — chimeric/supplementary | absent | *withdrawn* |
| YH229 | NC_030973.1 | 560,000 | 570,000 | 10,000 | PASS | unique | fail — chimeric/supplementary | absent | *withdrawn* |
| YH229 | NC_030978.1 | 70,000 | 80,000 | 10,000 | PASS | unique | fail — cerevisiae identity higher | absent | *withdrawn* |
| YH229 | NC_030980.1 | 560,000 | 588,913 | 28,913 | FAIL | recurs | non-decisive (mixed) | absent | *withdrawn* |
| YH229 | NC_030982.1 | 1,040,000 | 1,050,000 | 10,000 | FAIL | recurs | non-decisive (margin 29.9) | absent | *withdrawn* |
| YH229 | NC_030983.1 | 10,000 | 20,000 | 10,000 | FAIL | unique | non-decisive (margin 2139.3) | absent | *withdrawn* |
| YH229 | NC_030985.1 | 0 | 10,000 | 10,000 | FAIL | recurs | non-decisive (margin 155.8) | absent | *withdrawn* |
| YH229 | NC_030985.1 | 740,000 | 747,934 | 7,934 | FAIL | recurs | non-decisive (margin 991.7) | absent | *withdrawn* |

† FAIL = block does not survive terminus masking (subtelomeric); PASS = survives, tested further.

‡ Unique = does not recur at the same coordinates in unrelated controls; recurs = fails.

§ Read-identity margin is retained for completeness but is circular by construction (reads were already selected by competitive mapping); not evaluable = masking/recurrence already excluded the block; non-decisive = margin computed but not treated as discriminating.

**Table S2. Sixteen re-analyzed panel isolates.** ((Avelar-Rivas, et al. 2026); BioProject PRJNA1138754).

| **Isolate** | **SRA accession** | **Reported introgression blocks** | **Clade** | **Group** |
| --- | --- | --- | --- | --- |
| YMX002071 | SRR30037137 | 9 | Wine/European | Six lowest-count |
| L022* | SRR30037022 | 12 | Mixed Origin | Six lowest-count |
| YMX005648 | SRR30037010 | 13 | Mixed Origin | Six lowest-count |
| MOC1M* | SRR30037020 | 18 | Wild North American | Six lowest-count |
| YMX001945 | SRR30037167 | 40 | Wild North American | Six lowest-count |
| 1103* | SRR30037079 | 51 | Tequila Distillery | Six lowest-count |
| YMX005570 | SRR30037107 | 99 | Mexican Agave 2 | — |
| YMX003697 | SRR30037211 | 121 | Mexican Agave 2 | — |
| YMX001922 | SRR30037034 | 143 | Mexican Agave 2 | — |
| YMX005588 | SRR30037093 | 160 | Mexican Agave 1 | MA1 clade |
| DL5* | SRR30037213 | 166 | Mexican Agave 2 | — |
| YMX003402 | SRR30037173 | 171 | Mexican Agave 1 | MA1 clade |
| YMX009824 | SRR30037004 | 258 | Mexican Agave 1 | MA1 clade |
| YMX009823 | SRR30037005 | 259 | Mexican Agave 1 | MA1 clade |
| YMX009826 | SRR30037002 | 267 | Mexican Agave 1 | MA1 clade |
| YMX009795 | SRR30037178 | 364 | Mexican Agave 1 | MA1 clade |

* Not YMX-named; ID as given in the source paper's own sample sheet.

**Table S3. Reference genome assemblies.**

| **Species / strain** | **Assembly accession** | **Notes** |
| --- | --- | --- |
| *S. cerevisiae S288C* | GCF_000146045.2 | Assembly R64 |
| *S. paradoxus CBS432* | GCF_002079055.1 |  |
| *S. paradoxus YPS138* | GCA_002079115.1 | Ten-species panel reference only; distinct from CBS432 |
| *S. eubayanus FM1318 / CBS12357ᵀ* | GCA_001298625.1 / GCF_001298625.1 (SEUB3.0) |  |
| *S. mikatae IFO1815* | GCF_947241705.1 |  |
| *S. kudriavzevii CR85* | GCA_900682695.1 |  |
| *S. jurei* | GCA_900290405.1 |  |
| *S. arboricola H-6* | GCF_000292725.1 |  |
| *S. uvarum CBS7001* | GCA_027557585.1 |  |
| *K. marxianus DMKU3-1042* | GCA_001417885.1 |  |
| *P. kudriavzevii CBS573* | GCA_003054445.1 |  |

**Table S4. Negative controls and worked example.**

| **Strain** | **Role** | **Accession** |
| --- | --- | --- |
| S288C | Short-read negative control | SRR2070491 |
| EM14S01-3B | Divergent-control worked example | ERR1309189 |
| DoF1 | ONT negative control | SRR39135318 |

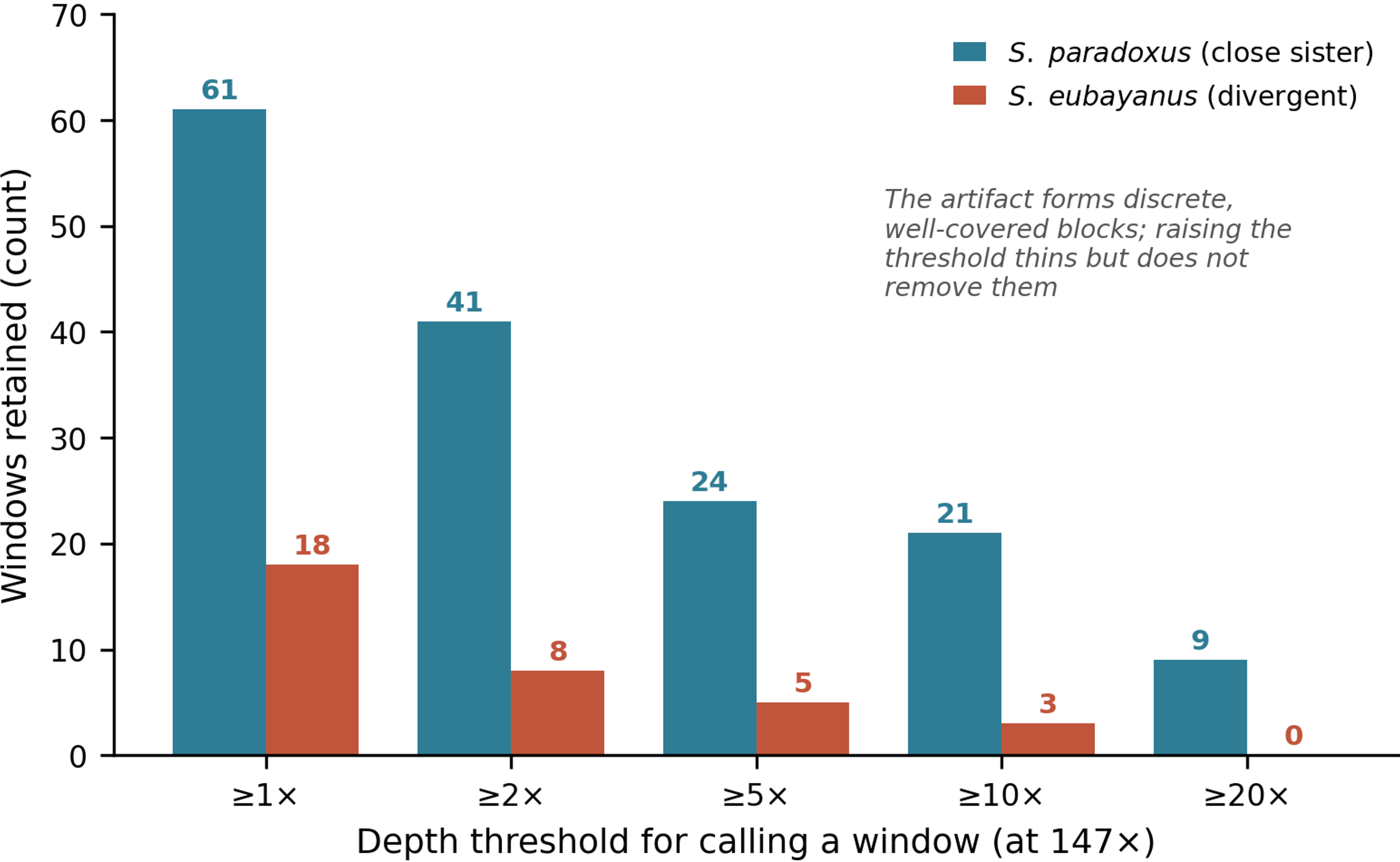

**Figure S1. Threshold sensitivity: window counts retained at ≥1×, ≥2×, ≥5×, ≥10×, ≥20× for each donor.** *S. paradoxus* and *S. eubayanus* window counts recovered from the S288C negative control at 147× (REF-naive), reclassified at successively stricter depth thresholds. *S. paradoxus* windows decline from 61 (≥1×) to 41, 24, 21, and 9 (≥20×). *S. eubayanus* declines in parallel from 18 to 8, 5, 3, and 0. The artifact forms discrete, well-covered blocks rather than diffuse noise. Raising the calling threshold thins the signal but does not remove it, and the divergent donor (*S. eubayanus*) is consistently smaller than the close sister (*S. paradoxus*) at every threshold.

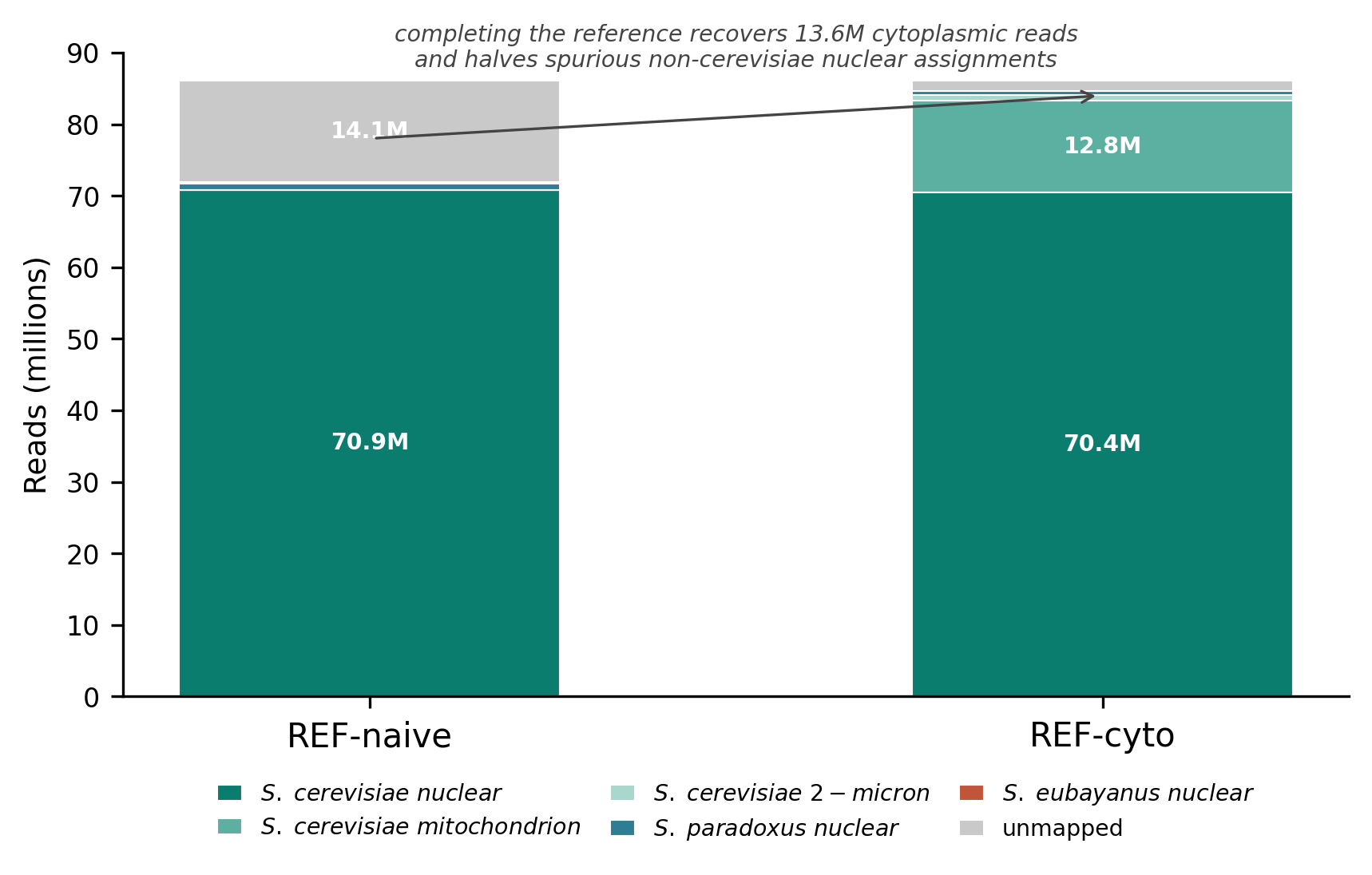

**Figure S2. Read-fate accounting for the short-read control: read counts by destination under each reference.** Full destination breakdown for the S288C control (SRR2070491) under REF-naive and REF-cyto. Completing the reference recovers 13.6 million reads previously left unmapped (12.81 million to the *S. cerevisiae* mtDNA and 0.82 million to the 2-micron plasmid) and reduces spurious non-*cerevisiae* nuclear assignments (*S. paradoxus*: 813,500 → 548,883 reads; *S. eubayanus*: 243,857 → 25,850 reads). Both totals reconcile exactly to the library's full read count (86.04-86.09 million).

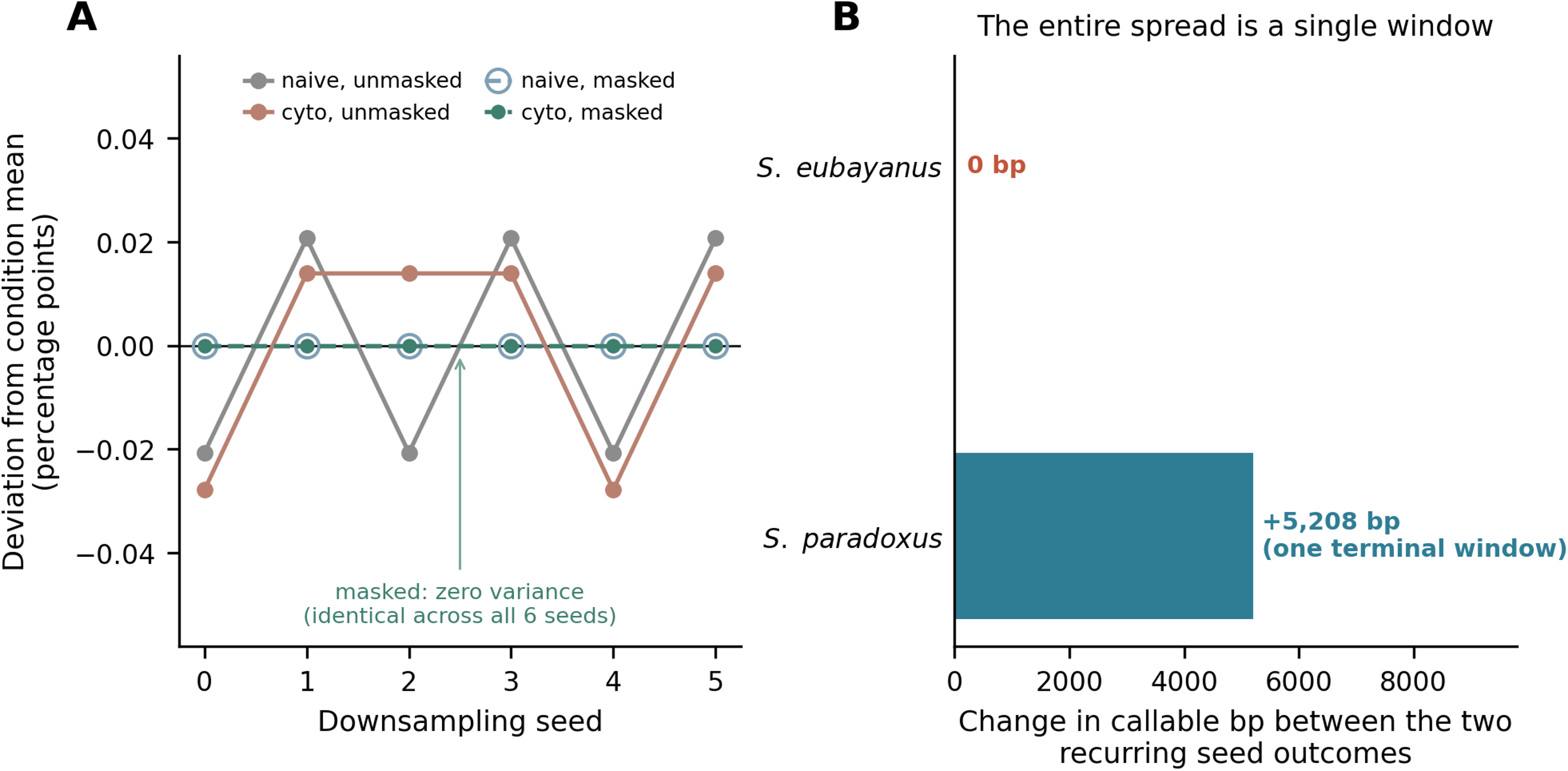

**Figure S3. The residual stochasticity of the floor is a single terminal window. (A)** Replicate down-sampling of the short-read negative control at 147×, six seeds, plotted as deviation from each condition's own mean so that all four conditions share one scale (means: REF-naive unmasked 2.038%, REF-cyto unmasked 1.646%, REF-naive masked 1.045%, REF-cyto masked 0.699%). The two unmasked conditions take exactly two distinct values across six seeds. Both masked conditions are identical across all six and are drawn with nested markers because they coincide exactly at zero deviation. **(B)** The difference between the two recurring unmasked outcomes is +5,208 bp of *S. paradoxus* and 0 bp of *S. eubayanus*, equal to the change in the callable denominator: a single ~5.2 kb terminal partial window crossing the ≥5× calling threshold.

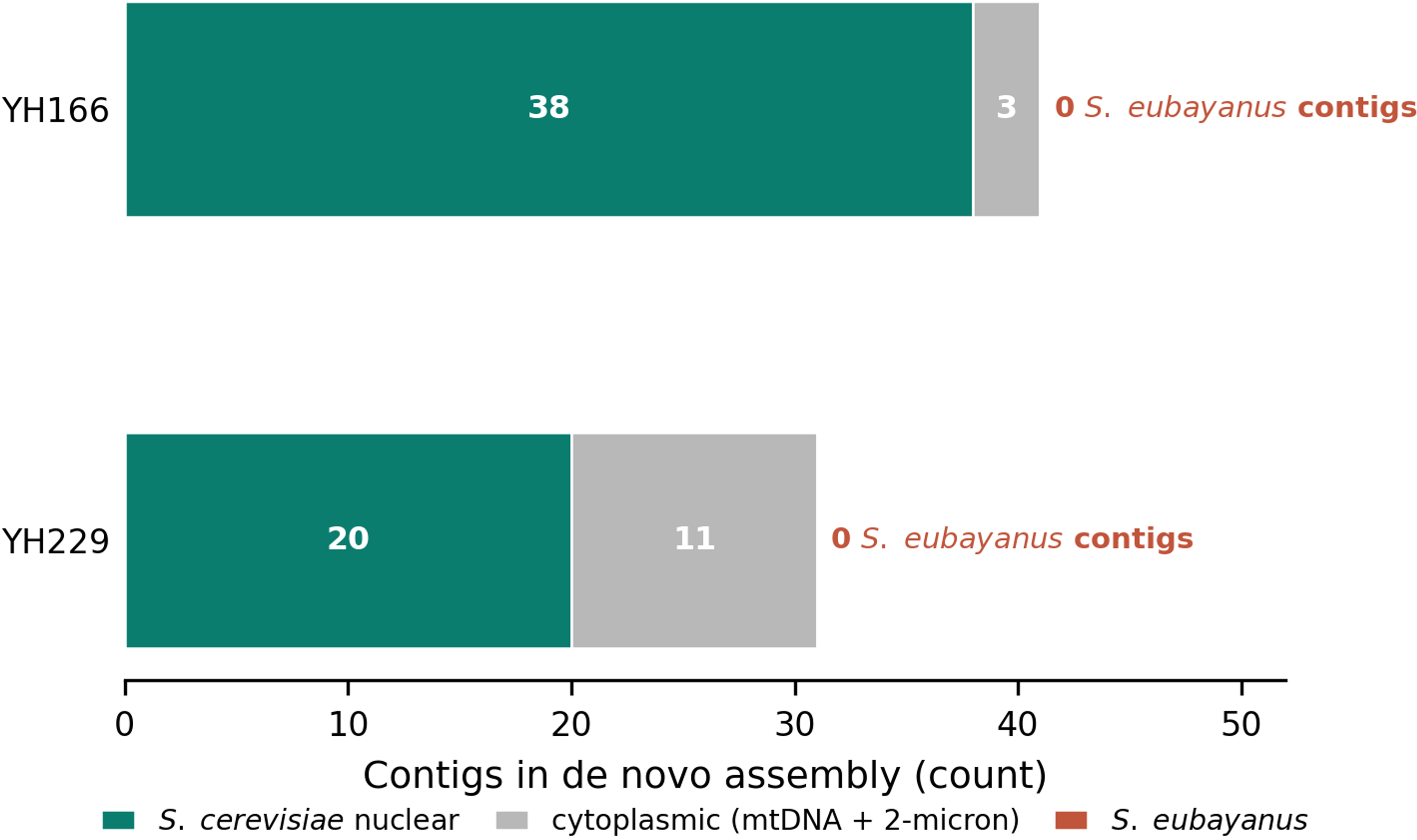

**Figure S4. Assembly cross-check: contig assignment by species for both withdrawn isolates.** *De novo* assembly contigs for YH166 (41 contigs; 12.40 Mb) and YH229 (31 contigs; 12.25 Mb), assigned by best alignment to the *S. cerevisiae* nuclear, cytoplasmic, or *S. eubayanus* reference. YH166 comprises 38 nuclear and 3 cytoplasmic contigs; YH229 comprises 20 nuclear and 11 cytoplasmic contigs. No contig in either assembly maps to *S. eubayanus*, consistent with the four-criteria withdrawal of both isolates' candidate introgression calls.
